# accuEnhancer: Accurate enhancer prediction by integration of multiple cell type data with deep learning

**DOI:** 10.1101/2020.11.10.375717

**Authors:** Yi-An Tung, Wen-Tse Yang, Tsung-Ting Hsieh, Yu-Chuan Chang, June-Tai Wu, Yen-Jen Oyang, Chien-Yu Chen

## Abstract

Enhancers are one class of the regulatory elements that have been shown to act as key components to assist promoters in modulating the gene expression in living cells. At present, the number of enhancers as well as their activities in different cell types are still largely unclear. Previous studies have shown that enhancer activities are associated with various functional data, such as histone modifications, sequence motifs, and chromatin accessibilities. In this study, we utilized DNase data to build a deep learning model for predicting the H3K27ac peaks as the active enhancers in a target cell type. We propose joint training of multiple cell types to boost the model performance in predicting the enhancer activities of an unstudied cell type. The results demonstrated that by incorporating more datasets across different cell types, the complex regulatory patterns could be captured by deep learning models and the prediction accuracy can be largely improved. The analyses conducted in this study demonstrated that the cell type-specific enhancer activity can be predicted by joint learning of multiple cell type data using only DNase data and the primitive sequences as the input features. This reveals the importance of cross-cell type learning, and the constructed model can be applied to investigate potential active enhancers of a novel cell type which does not have the H3K27ac modification data yet.

**Availability:** The accuEnhancer package can be freely accessed at: https://github.com/callsobing/accuEnhancer

## Introduction

Understanding the effect of gene regulation to human health is a fundamental scientific problem and emerges to be a critical task of translational genomics in the clinics. Due to the rapid growth of next-generation sequencing (NGS) technologies (Mardis 2008), exploring individual genomes much more thoroughly becomes possible than ever. By means of whole genome sequencing (WGS) (Cirulli and Goldstein 2010), we can now not only identify the variants located in the coding regions but also discover a massive number of variants located in the non-coding regions (Zhang and Lupski 2015). The functional and regulatory importance of the coding regions can be revealed by various annotations, such as protein function, pathway information, and other annotation resources from public databases. In contrast, the annotation of non-coding regions remains largely unknown (Griffiths-Jones, Moxon et al. 2005). Due to the recent developments of international consortiums like ENCODE (Consortium 2004), Roadmap (Bernstein, Stamatoyannopoulos et al. 2010), GTEx (Lonsdale, Thomas et al. 2013) and other major genomic research groups for their attempts to characterize the regulatory genomes by producing various types of experimental data in the past decade, these data have made perfect resources for investigating the regulatory grammar in the noncoding regions.

Despite a large number of accumulated experimental datasets, how to investigate the active regulatory elements in different cell types from these data is still an open question. Many studies implied that the non-coding regions can be divided into different categories, such as promoters, enhancers, insulators, silencers and other functional units (Ernst and Kellis 2012). While promoters can control the basic expression profiles of the downstream proximal genes, the enhancers can activate expression of multiple distal genes by chromosome looping or other mechanisms. These regulatory grammars are often considered as cell type specific processes (Spitz and Furlong 2012). The active enhancers within a cell type are genomic loci containing several transcription factor (TF) binding sites and generally located in the accessible chromatin regions (Andersson, Gebhard et al. 2014). Recent genome-wide investigation of the epigenetic marks has discovered that active enhancers are associated with certain histone modification signals, such as H3K4me3 (Chen, Chen et al. 2015), H3k27ac (Heintzman, Stuart et al. 2007, Creyghton, Cheng et al. 2010), and H3k9ac (Karmodiya, Krebs et al. 2012), and these histone modification data make valuable sources for predicting active enhancers. There are several high throughput experimental approaches to define the active enhancers within each cell type. The first is to map the binding sites of the TFs to the genome using ChIP-seq data. This approach requires not only a large amount of ChIP-seq experimental data of TF binding but also the knowledge of the proper TF combinations (Shlyueva, Stampfel et al. 2014). The second approach is to identify the binding profiles of the histone acetyltransferase EP300, a TF which recruits cofactors to initiate the transcription activation process by acetylating chromatin (Lee, Karchin et al. 2011). The third approach is to use the DNase-I hypersensitivity sites as the potential active enhancers (Wilken, Brzezinski et al. 2015). However, the open chromatin regions consist of many types of regulators, including promoters, enhancers, silencers, and insulators, using DNase-I hypersensitivity sites as the potential active enhancers lacks the specificity to identify active enhancers precisely. While enhancer elements are known to be associated with certain histone modifications, another approach is to use epigenetic marks, such as H3K27ac, to identify the active enhancers (Creyghton, Cheng et al. 2010). The histone modification data, H3K27ac, has been used to distinguish active from inactive enhancers in previous studies (Creyghton, Cheng et al. 2010, Natoli and Andrau 2012, Zhu, Sun et al. 2013).

In addition to the experimental methods mentioned above, there are several computational approaches that have been proposed in the past decade. The problem can be tackled from different aspects, such as classification of the chromatin states (ChromaSig (Hon, Ren et al. 2008), ChromHMM (Ernst and Kellis 2012), Segway (Hoffman, Buske et al. 2012)), prediction of the p300 binding sites (CSIANN (Firpi, Ucar et al. 2010), RFECS (Rajagopal, Xie et al. 2013), DELTA (Lu, Qu et al. 2015), REPTILE (He, Gorkin et al. 2017)), prediction of the H3k27ac peaks (PEDLA (Liu, Li et al. 2016), EP-DNN (Kim, Harwani et al. 2016)), and prediction of the experimentally validated enhancers from VISTA (Visel, Minovitsky et al. 2007) or FANTOM (Bono, Kasukawa et al. 2002) databases (DEEP (Kleftogiannis, Kalnis et al. 2015), EnhancerDBN (Bu, Gan et al. 2017), and BiRen (Yang, Liu et al. 2017)). Among all the developed models, only a handful of studies attempted to predict enhancer activities across cell types (using the data of one cell type as the training data to predict the enhancer activity of another cell type), which is still a difficult task when compared to the within-cell type predictions (using partial data of one cell type as the training data to predict the enhancer activity of the holdout data from the same cell type).

In this regard, the objective of this work is to build a cross-cell type model for active enhancer predictions. We first constructed a within-cell type predicting model by using sequence information and several epigenetics marks (H3K27me3, H3K36me3, H3K4me1, H3K4me2, H3K4me3, H3K9ac, and DNase) to evaluate the feasibility of using convolutional neural networks (CNN) (Lawrence, Giles et al. 1997) in enhancer prediction. When a comprehensive set of functional data is used as the input features, not only the within-cell type models deliver satisfied performance, but also the cross-cell type models. However, this means all functional data features must be available before the prediction of active enhancers for a novel cell type can be made. To reduce the cost of predictions, we aim to reduce the types of functional data used in constructing the prediction models. The DNase data retrieved from DNase-I hypersensitivity sites is considered because it is ubiquitously available across different cell types and highly related with the active enhancers. In this study, we develop accuEnhancer that can accurately predict cross-cell type enhancer activity by using only the DNase data along with the genomic sequence. First, we enlarge the range of the functional data explored when constructing the models. We then collect data from multiple cell types (50 in total) and use them as the input data of accuEnhancer to demonstrate the power of the developed joint training framework. The main concept of the joint training using multiple cell types is to take the advantage from the variability between different cell types, in order to discover critical hidden patterns embedded in active enhancers. We further utilize the pre-trained filter weights constructed from the DeepC (Schwessinger, Gosden et al. 2020) study, which trained the deep neural nets with hundreds of epigenomics data, as the initial weights in the genomic sequence module of accuEnhancer to improve the model performance. All the three distinct features of accuEnhancer, enlarging the explored region of DNase, joint training of multiple cell types, and using pre-trained weights from DeepC, are shown to provide promising improvements on the prediction performance. In the end of this study, we tested accuEnhancer on experimentally validated enhancers in the VISTA database and compared the performance of accuEnhancer with other existing tools. In summary, accuEnhancer is the first methodology that attempts to integrate multiple cell types as the training dataset for predicting enhancer activity using only the DNase data along with genomic sequences. The integration of multiple cell type data with joint learning makes accuEnhancer have the superior model performances and provides different aspects to tackle the enhancer predicting problems. The accuEnhancer package and all the preprocessing scripts can be accessed on the GitHub repository (https://github.com/callsobing/accuEnhancer).

## Results

### Network structure of accuEnhancer

The proposed method, accuEnhancer leverages the characteristics of CNN, a feedforward neural network with a large number of neurons widely used in image processing, speech recognition, and natural language processing, to build the model of predicting active enhancers. The architecture of accuEnhancer consists of two different modules (Figure 1), the *genomic sequence module* and the *functional data module*. The numbers of filters in both genomic sequence and functional data modules were set to 64 with a length of 16. The final setting was determined after a series of evaluations across different parameter settings. The stride in the max-pooling layer is set to 1, and the fully connected layers both have 32 hidden nodes. The sequence length of each instance is set to 200 base pairs (bps) and the sequences will be further one-hot encoded into a 4×200 matrix for downstream analysis. Since the DNase data (DNase-I hypersensitivity sites) is a region-based functional data, we attempted to expand the observed DNase region from [-100 bps, +100 bps] to [-2500 bps, +2500 bps] (the center of the input sequence is set to zero as the reference point) and divide this region into 25 bins, of which each has 200 bps in length. This will result in a 1×25 matrix that is used as the input for the functional data module.

**Figure 1.**
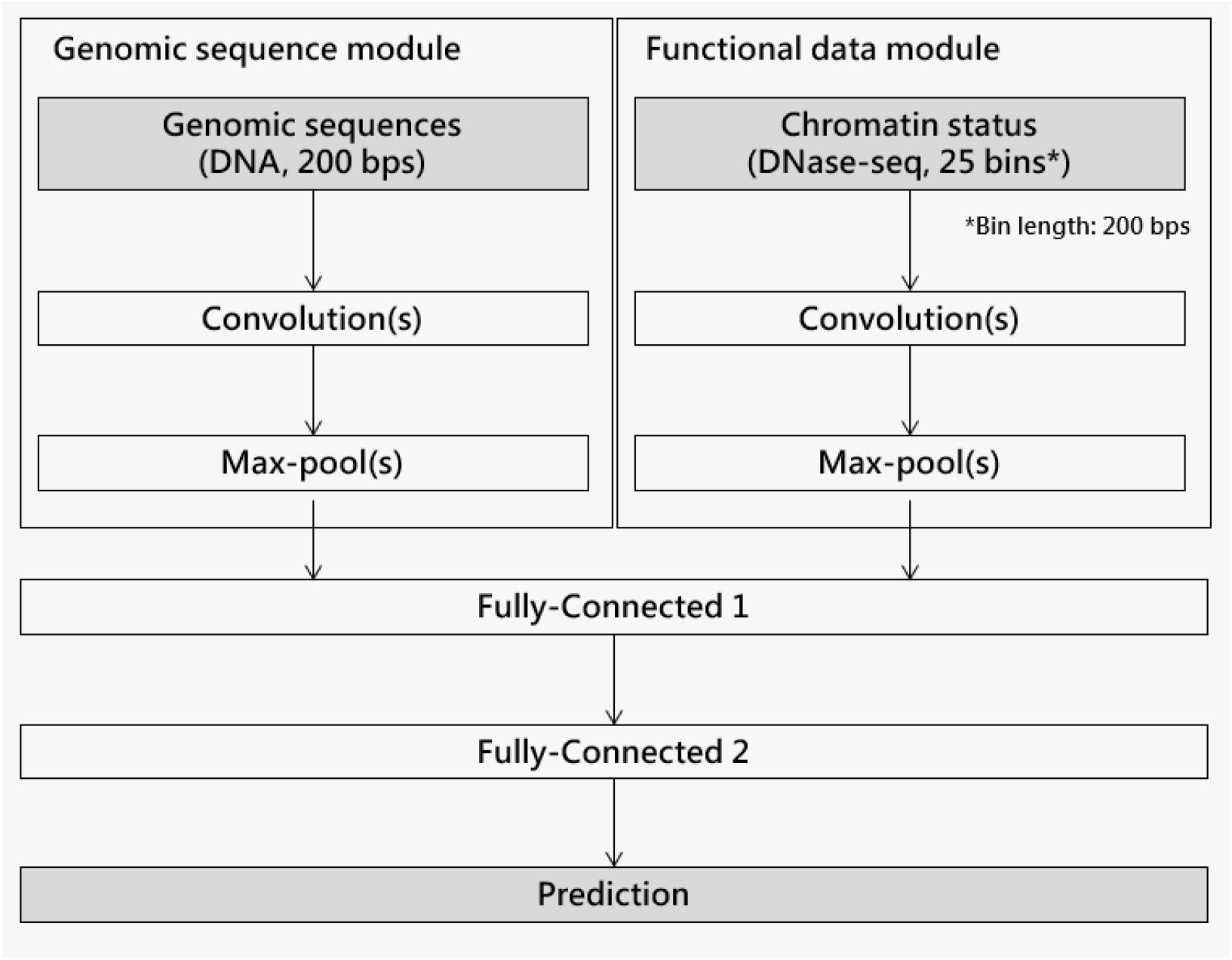
The network structure of accuEnhancer. The framework consists of two modules, the *genomic sequence* module and the *functional data* module. The two modules learn within module patterns locally and interact with another module in the fully connected layers.

A previous work, DeepC, used 936 chromatin-accessibility and epigenetic datasets to train a deep CNN to predict chromatin accessibility based on DNA sequence across cell types. The DeepC network architecture was adapted from DeepSEA, a deep learning-based method for predicting the chromatin effects. DeepC used five convolutional layers with ReLU as the activation function, followed by 1D max pooling and dropout at every convolutional layer. The DeepC model learned to recognize chromatin features from DNA sequences first and then combined the underlying sequence patterns to predict chromatin interactions. The genomic sequence module of accuEnhancer leveraged the DeepC-pretrained weights to predict the chromatin features from the given DNA sequences and replaced the final output layer of DeepC with two convolution layers for further training. In this study, we tested the within-cell type model performances with and without using the pre-trained weights from DeepC, respectively. The result shows that with the pre-trained weights from DeepC, the model achieved better F1 score on the testing dataset (Supplementary information: Figure S1). Therefore, the pre-trained weights are further used throughout the rest analyses of this study. Figure S1 also provided us with the good performance that a within-cell type model can deliver when using a comprehensive set of functional data as the input features.

### Integrating data from multiple cell types improves enhancer prediction accuracy

In order to build models to predict cross-cell type enhancer activity, we first attempted to select proper functional data as the input features of the deep neural networks. Although combining various kinds of functional data can predict active enhancers accurately across cell types (Supplementary information: Table S1), the cost of conducting the ChIP-seq experiments of all histone markers makes it barely impossible to generate all the required data. DNase data is one of the functional data type that has been reported as highly related to active enhancers and the DNase data is also one of the largest collections of functional data in the ENCODE database. In this regard, accuEnhancer combines genomic sequences and DNase information related to the given loci to construct the predicting models. We used the model trained from one single cell type (H1-hESC) to predict another cell type (HepG2). The performance (F1 score: 0.29) is poor (Figure 2), and the area under ROC is 0.783. Since the genomic context between different cell types may vary a lot, the main concept of accuEnhancer is to learn the complex regulatory grammars from different cell types of the training data. For this purpose, we selected 50 cell types with publicly accessible DNase and H3K27ac data as our training data, where the H3K27ac data is used to label each of the training instances as positive or negative. The correlation of active enhancers between these 50 cell types are calculated based on the H3K27ac ChIP-seq peaks of each cell type (Supplementary information: Figure S2). The analysis showed that the similarity of active enhancers between these 50 cell types is not high, which is good for providing active enhancers from multiple cell types for the training purpose. We incrementally increased the number of the integrated cell types to build the accuEnhancer models and tested the performances on the independent cell type (HepG2). The performances of the models integrating 1, 3, 5, 9, 20, 30, 50 cell types are shown in Figure 2. The ROC curves showed that the models constantly improved performances while including more cell types as the training data. The final model that merges 50 cell types achieved area under ROC (AUC) of 0.958, which is satisfied for predicting active enhancers in novel cell types for future applications.

**Figure 2.**
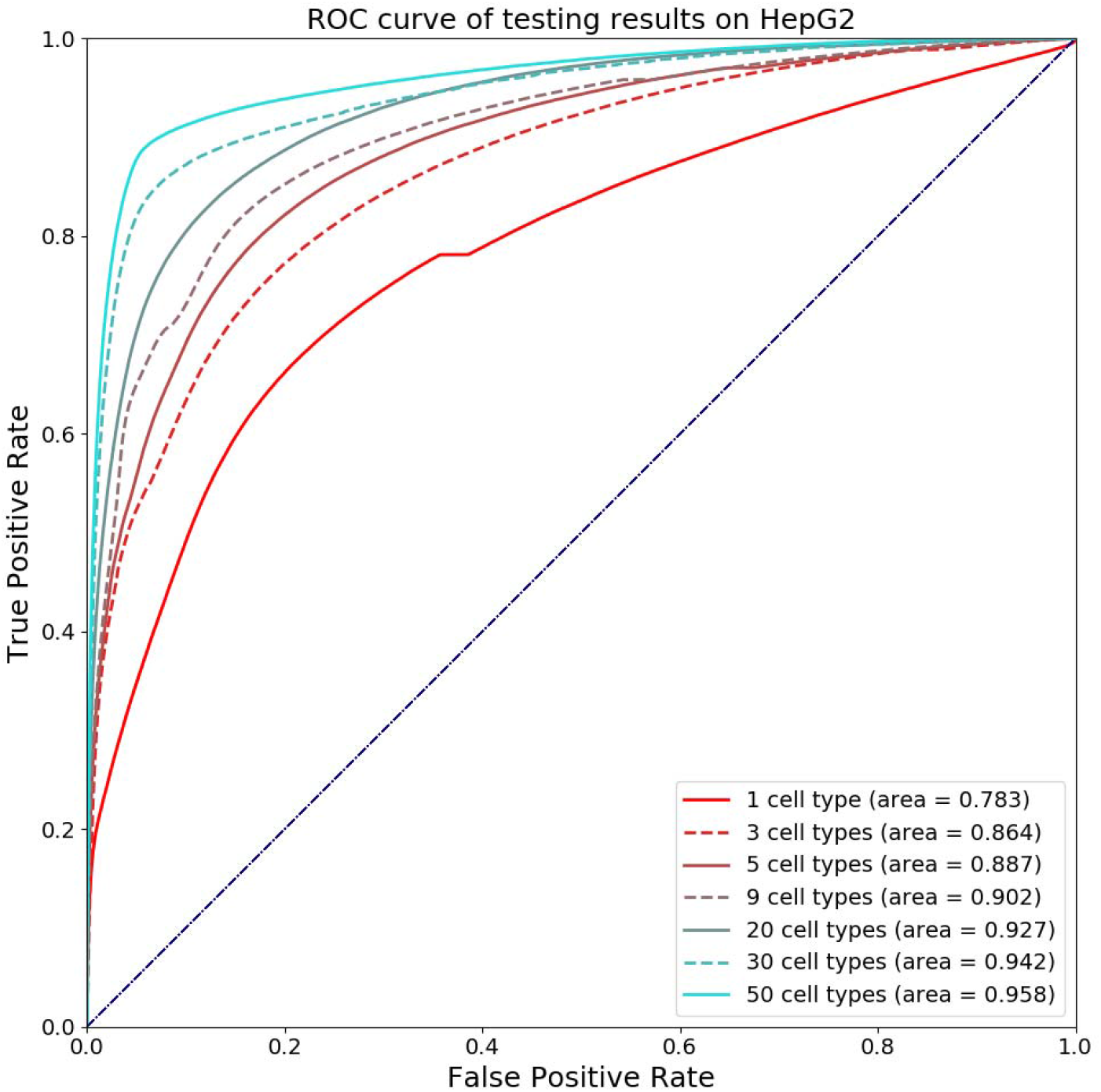
The ROC curves of accuEnhancer models with different numbers of cell types included in the training data. The result indicates that the models constantly improved the performances while including more cell types in the training data.

### Integrating data from multiple cell types helps to identify novel active enhancers

Since the AUC of the accuEnhancer model increased apparently as we integrated more training cell types, we would like to investigate if the better performance came from the enhancers that have been previously observed in the training dataset. We first listed all the loci of 200 bps bins of active enhancers in the 50 training cell types and in the HepG2 cell type, respectively. Next, we identified the enhancers which are active in the HepG2 but never active in any of the 50 cell types used in training. This resulted in a total of 24,936 bins of HepG2 novel enhancers. The model prediction outcomes of these 24,936 HepG2 novel enhancers are shown in Figure 3. The result revealed that when an active enhancer is not present (novel) in the training dataset, only 1,304 enhancers can be predicted as active enhancers in the single cell type model. On the other hand, 10,705 enhancers can be correctly identified as active enhancers in the model combining 50 cell types. The results suggest that integrating data from multiple cell types can not only increase the model performance by observing more active enhancers from other cell types, but also detecting some hidden patterns to identify active enhancers that have never been observed as active enhancers from the previous cell types. We also compared the relations between recall and precision of different models in predicting HepG2 active enhancers (Figure 4). The recall rates increased from 0.18 to 0.83, and the precision rates slightly increased from 0.64 to 0.83, when number of cell types integrated in accuEnhancer increased. As the recall rate of the HepG2 novel active enhancers is concerned (Supplementary information: Figure S3), not only the recall rate of ‘non-unique’ enhancers increases when the number of cell types included in the training process increases, but also the recall rate of ‘novel’ enhancers. This demonstrates the ability of accuEnhancer in retrieving complicated regulatory patterns through the diversity of active enhancers from multiple cell types.

**Figure 3.**
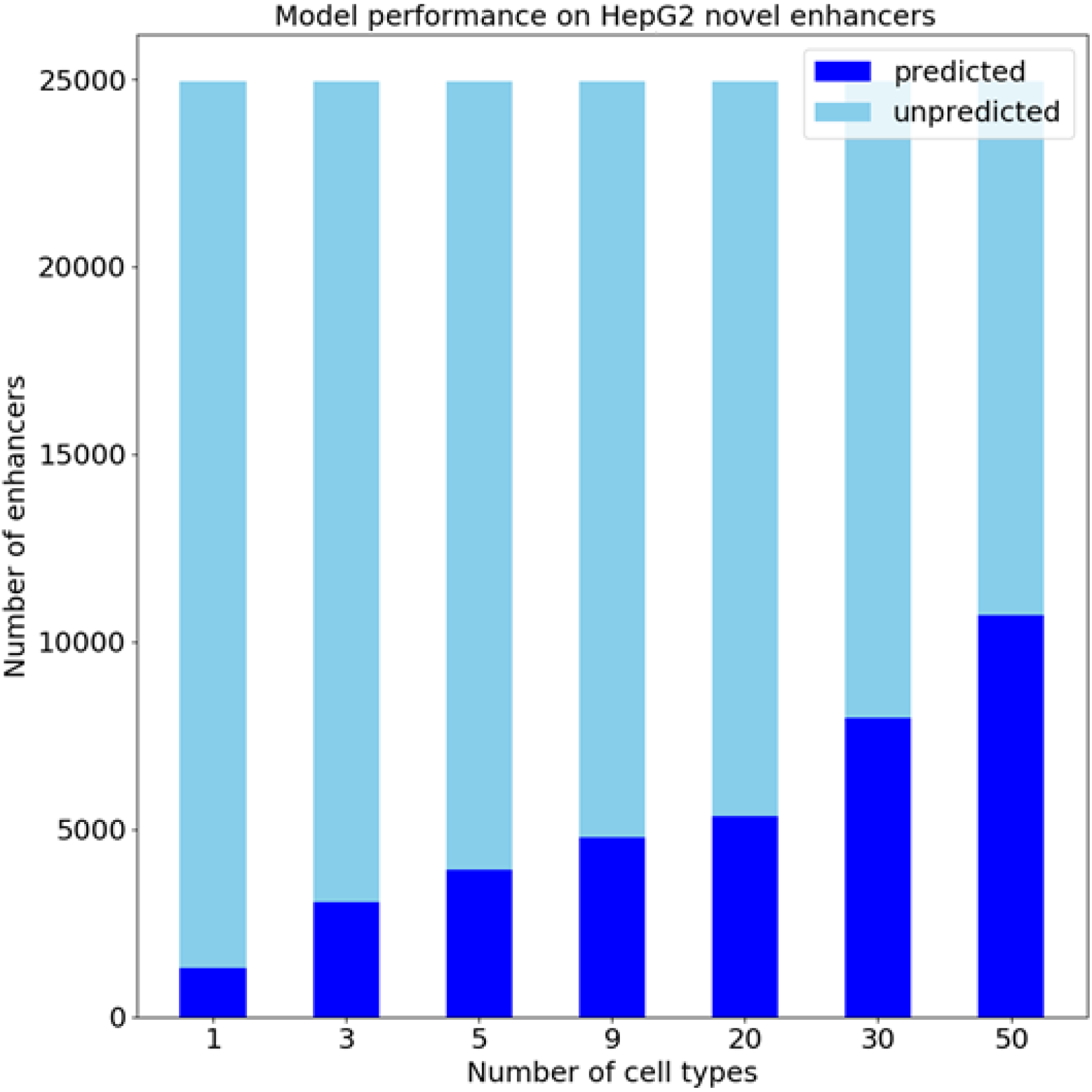
The performances of accuEnhancer in predicting active enhancers uniquely existing in the testing cell type (HepG2), called HepG2 novel enhancers. Even these enhancers have never been labeled as active enhancers in the training data, accuEnhancer still can correctly predict some of them as active enhancers in HepG2. The result suggests that accuEnhancer not only recognizes potential active enhancers through previously reported enhancers, but also has learned some hidden patterns to identify the active enhancers which have not been labeled active enhancers in the training data.

**Figure 4.**
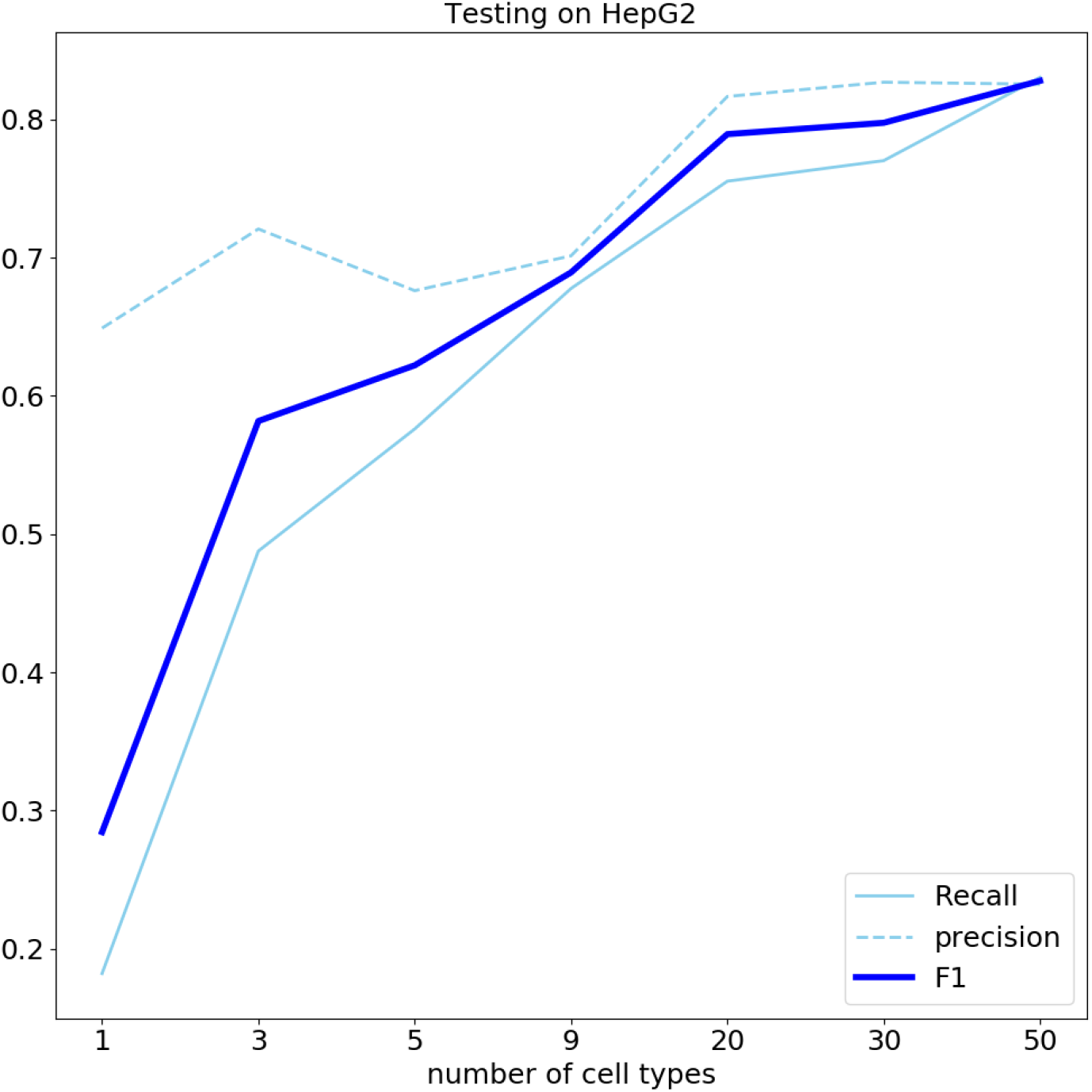
The comparison of the recall rates, the precision rates, the F1 scores of different models in predicting HepG2 active enhancers. The result indicates that the model tends to make more predictions on active enhancers, when integrating more cell types, but not sacrifices the precision.

### accuEnhancer outperforms other algorithms for predicting VISTA enhancers

accuEnhancer demonstrated the ability to predict cross-cell type active enhancers by integrating data from multiple cell types in human. Here, we further tested whether the strategy to integrate multiple lines of data to predict active enhancers also works in other species. For this purpose, we collected the active enhancers from the VISTA database, a central resource for experimentally validated human and mouse noncoding fragments with gene enhancer activity as assessed in transgenic mice, and constructed accuEnhancer models in mouse. We collected the active enhancers data from mouse embryo tissues, including hindbrain, forebrain, neural tube, and midbrain, at embryonic day 11.5 with available DNase ChIP-seq data. We used these four tissues in this study to train four accuEnhancer models separately, each time using three tissues as training data and the rest one tissue as testing data. To compare with other algorithms, including CISANN, DELTA, REPTILE and RFECS, we also trained accuEnhancer on mouse embryonic stem cell (mESC) as REPTILE suggested. The predicting results for the four tissues are shown in Figure 5. accuEnhancer shows slightly better predicting AUPRC than existing methods based on the mESC training data, while the performance of joint training data of other three cell types (denoted as ‘3 tissues’) outperformed all other methods with significant differences. We further examined the area under ROC (AUC) for hindbrain and discovered that accuEnhancer trained on mESC has a very similar AUC (0.793) with REPTILE (0.725), but the AUC of the accuEnhancer increased to 0.981 as the model uses other three tissues as the training data (Figure 6). This result not only shows accuEnhancer works in human, but also shows superior performances in mouse tissues. This suggested that when with DNase data, accuEnhancer can predict active enhancers in novel cell types within same species with high accuracy. accuEnhancer outperforms the previous works in predicting VISTA enhancers, which demonstrates that the strategy to integrate multiple cell types for joint training can boost the model performance in predicting active enhancers.

**Figure 5.**
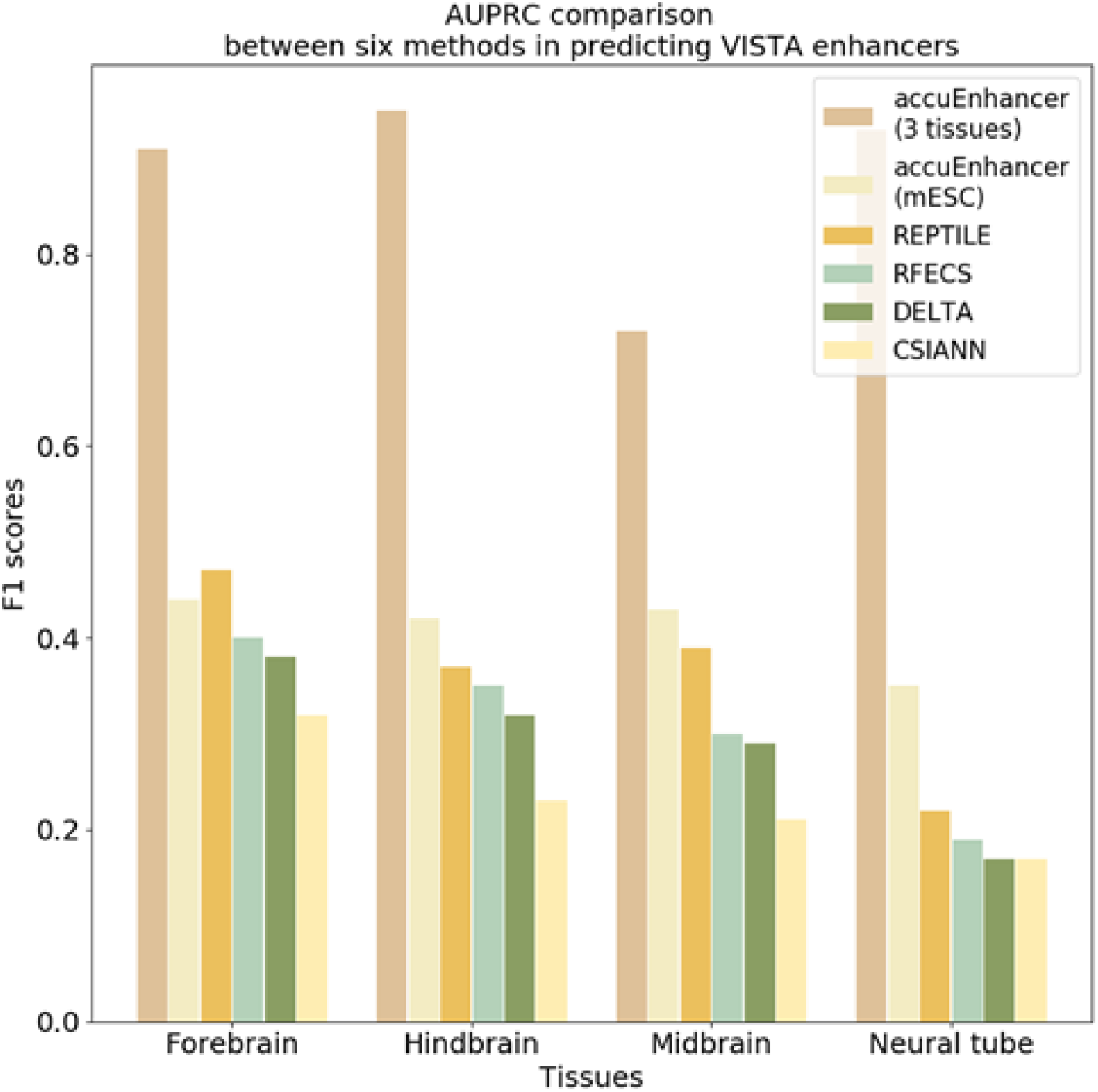
Performance evaluation between accuEnhancer and four existing enhancer predicting tools based on VISTA enhancers. In this analysis, we picked four tissues with available DNase experimental data. For each run of the four experiments of accuEnhancer (denoted as ‘3 tissues’), we used three tissues to train accuEnhancer and leave one tissue out for the testing purpose.

**Figure 6.**
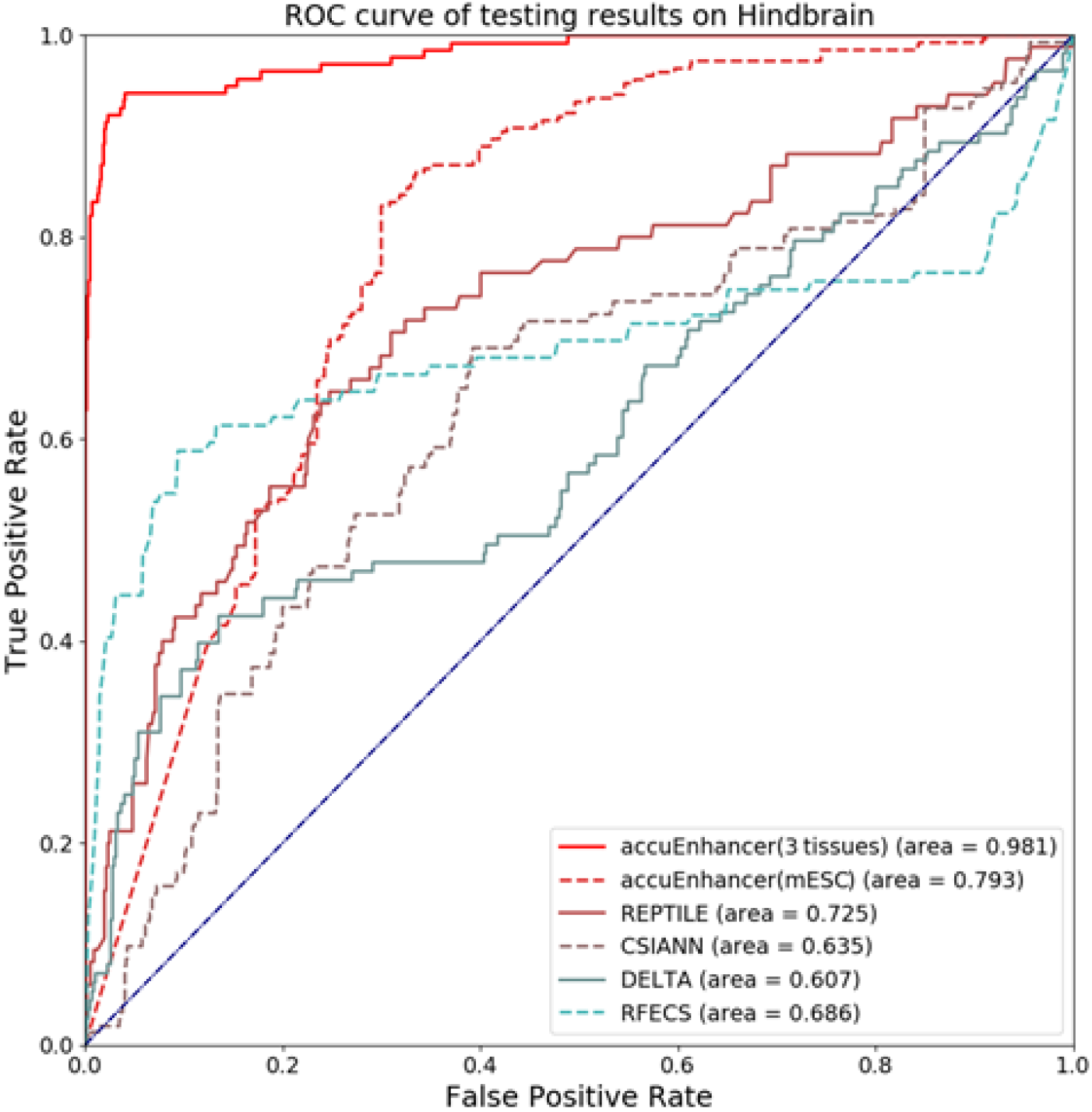
The ROC curves of the prediction results of accuEnhancer and other four studies on VISTA enhancers in Hindbrain. This result shows accuEnhancer not only works in human, but also shows superior performance in mouse tissues. This suggested that accuEnhancer can predict active enhancers across cell types with high accuracy by using only DNase data along with the genomic sequences.

## Discussions

In this study, we demonstrated the effectiveness of deep learning framework and revealed the potential of using only the DNase-derived features along with the sequence information to construct the model for predicting active enhancers across different cell types. accuEnhancer also indicates that the model performances can be largely enhanced by integrating data from different cell types. The successful application of accuEnhancer on predicting the VISTA enhancers in different tissues also showed that the model can apply to other species and identify active enhancers based on the DNase-seq and genomic sequence data only. Despite there are many databases and consortiums working hard to generate functional data from different cell lines, tissues, or primary cells, many cell types still lack sufficient data to annotate the location of active enhancers along the genome. The construction of accuEnhancer aims to locate the active enhancers within a novel cell type and requires only limited functional data, the DNase-seq.

As the number of DNase bins used in the functional data module is concerned, we tested different numbers of DNase bins adopted the functional data module (Supplementary information: Figure S5) when constructing the 50-cell type model. The results demonstrated that using 13 bins largely improved the prediction performance, while using 25 bins provided stable predictions on the HepG2 testing data.

As the resource usage in constructing the models is concerned, we investigated the memory usage and the training time of constructing models including different numbers of cell types (Supplementary information: Figure S4). The hardware specification is as follows: CPU: Intel(R) Xeon(R) Gold 6126 CPU@2.60GHz; GPU: geforce-rtx-2080-ti; Memory: 360 GB. Figure S4 reveals the demanding of hardware increases when the number of cell types used in joint training increases. This reveals the importance of sharing accuEnhancer as a public resource for enhancer activity prediction.

New experimental techniques like ATAC-seq^35^, which is faster and much more sensitive than DNase-seq, might be a better source to interrogate chromatin accessibility. Since data acquired from DNase-seq are very sparse, the ATAC-seq with better resolutions might help the model to perform better on specific scenarios. In the future, we will investigate whether ATAC-seq can be used as a replacement of DNase-seq and take advantage of the growing number of ATAC-seq datasets.

## Methods

### Input data and features

When invoking CNN to build the prediction models, we prepared each training instance with a 200-bp sequence. We used the human genome version GRCh38 as the reference and split the entire genome into non-overlapped bins with 200 bps in length. Then, we encoded the DNA sequences from a string consisted of four bases (A,C,G and T) into a matrix with four channels using the one-hot encoding method. The one-hot encoding uses one 1 and three 0s to represent each nucleotide. For example, A, C, G and T will be encoded as (1000), (0100), (0010) and (0001) respectively. Therefore, each training instance would be a 4×200 matrix to represent the sequence feature, where each row representing a specific locus and each column representing a particular nucleotide, after applying the one-hot encoding method. Other than genomic sequences, previous studies have shown that functional data, such as histone modification and chromatin accessibility information, is in correlation with the active enhancers. In this study, we selected DNase-I hypersensitive sites, which correspond to the openness of the chromatin structure, as the functional feature to train the prediction models. The narrow peak files of each cell type-specific DNase data were downloaded from the ENCODE database. In order to capture the broader conformation of the chromatin structure around the bin to be predicted, we expanded the feature extraction region to −1,200 bps and +1,200 bps around the mid-point of the target bin. The extended regions are further separated into 24 bins with 200 bps in length and the scores for each bin are the highest signal value in the narrow peak file that overlapped with it (Supplementary information: Figure S6).

### Construction of the positive and negative training data

Several enhancer markers have been proposed during the last few decades, including EP300 binding sites, H3K27ac binding profiles, DNase hypersensitivity sites, and eRNA transcribed regions. Among these experimental data, H3K27ac and EP300 are the most widely used enhancer markers. The ENCODE database has only 96 EP300 experimental data, but contains 443 H3K27ac experimental data as public resources. While these two marks both show high correlation with active enhancers, we selected the h3k27ac ChIP-seq data, which is much available than EP300, to label the instances for training and testing. To build a cross-cell type mode, we selected 50 cell types (Supplementary information: Table S2) from ENCODE database. And we downloaded the narrow peak files called by the standard ENCODE pipeline as our raw training data. To align data from different cell types, we used a sliding window of 200 bps to divide the whole genome into bins. These 200 bps bins will further intersect with the called H3K27ac peaks, and the overlapped bins will be labeled as positive instances. While the negative data also plays an important role for a classification problem, we picked random genomic sequences with a length of 200 bps (10 times of the number of positive bins) as the negative instances. In order to test the performance of the prediction model combining multiple cell types, we simply merged the training data from different cell types and removed redundant training instances by keeping only one instance among those with the same sequences as the merged training data.

### Testing data and performance assessment

To evaluate the model performance of accuEnhancer, we selected HepG2 as the independent testing data. The model performance is evaluated using the AUC (area under ROC curve). The ROC curve is a graph showing the performance of a classification model at all classification thresholds, which consists of TPR (true positive rate) and FPR (false positive rate). AUC provides an aggregate evaluation of performances across all possible classification thresholds. Other metric, such as F1 score, recall, precision, are also used in this study.

### Performance comparison with recent studies on VISTA validated enhancers

REPTILE, a method to locate enhancers based on genome-wide DNA methylation and histone modification profiling, has compared its performance with other three methods on the VISTA validated enhancer dataset. We downloaded the VISTA validated enhancers and the prediction outputs of the four methods from the REPTILE GitHub page (https://github.com/yupenghe/REPTILE). We trained our model by using the VISTA validated enhancer sequences and the mESC DNase data (https://www.encodeproject.org/experiments/ENCSR319PWR/) using the same data preprocessing procedure as previously described. We also downloaded the testing DNase data from four tissues within the mouse E11.5 life stage, including forebrain (https://www.encodeproject.org/files/ENCFF090SFC/@@download/), midbrain (https://www.encodeproject.org/files/ENCFF400KSK/@@download/), hindbrain (https://www.encodeproject.org/files/ENCFF518OYM/@@download/), and neural tube (https://www.encodeproject.org/files/ENCFF002XOR/@@download/).

## Code availability

The codes used to prepare the input data, construct the models, and make predictions are available at the GitHub page: https://github.com/callsobing/accuEnhancer The 50-cell type model is also available at the above GitHub page.

## Acknowledgement

We would like to thank the ENCODE Project and VISTA enhancer browser for making their data publicly available. We appreciate Ko-Han Lee and Ting-Fu Chen for their valuable comments and suggestions.

## Author information

Y.-A.T. and C.-Y.C initiated and designed the study. Y-A.T. developed the analysis procedures, collected and organized the data, performed the analysis, and wrote the manuscript. C.-Y.C. guided the research direction and provided ideas to improve the methods. W.-T.Y. and T.-T.H. tested the codes, verified the analyses, and helped the release of the codes. J.-T.W., Y.-C.C., and Y.-J.O. helped in experimental design and gave useful comments. All authors read and approved the final manuscript.

## Supplementary information

**Figure S1.**
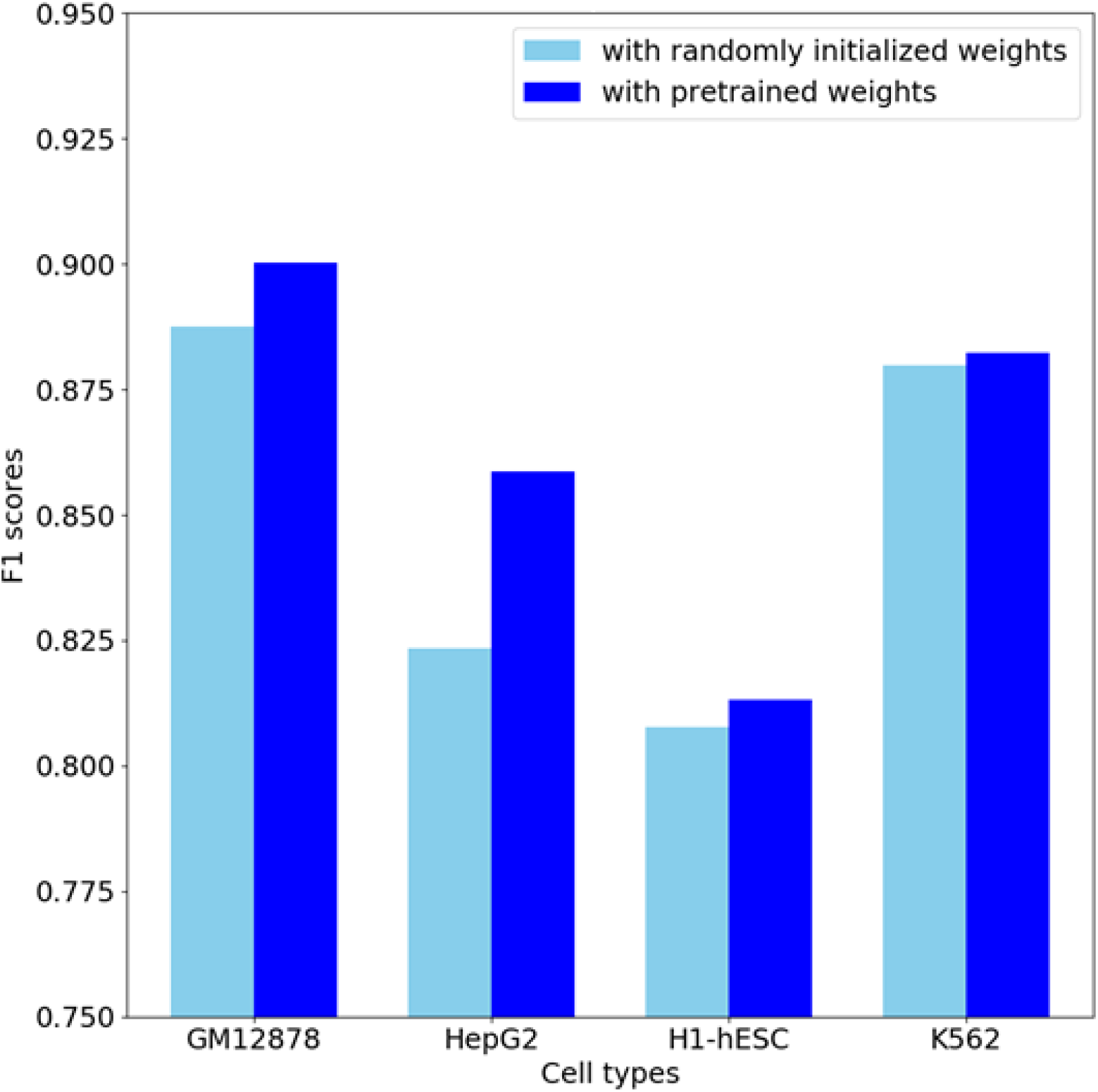
The result of the models trained by initializing the filter weights of the genomic sequence module with and without the pre-trained weights from DeepC. The validation set is a held-out validation set of the original training data. The ratio between training data and validation data is 10:1. We observed that even the training F1 scores are comparable between models with and without the pre-trained weights, the models trained with pre-trained weights can provide more robust results than the models trained with randomly initialized weights. This model uses H3K27me3, H3K36me3, H3K4me1, H3K4me2, H3K4me3, H3k9ac, DNase as the input features, along with the genomic sequences, in contrast to accuEnhancer that uses only DNase data in the functional data module.

**Figure S2.**
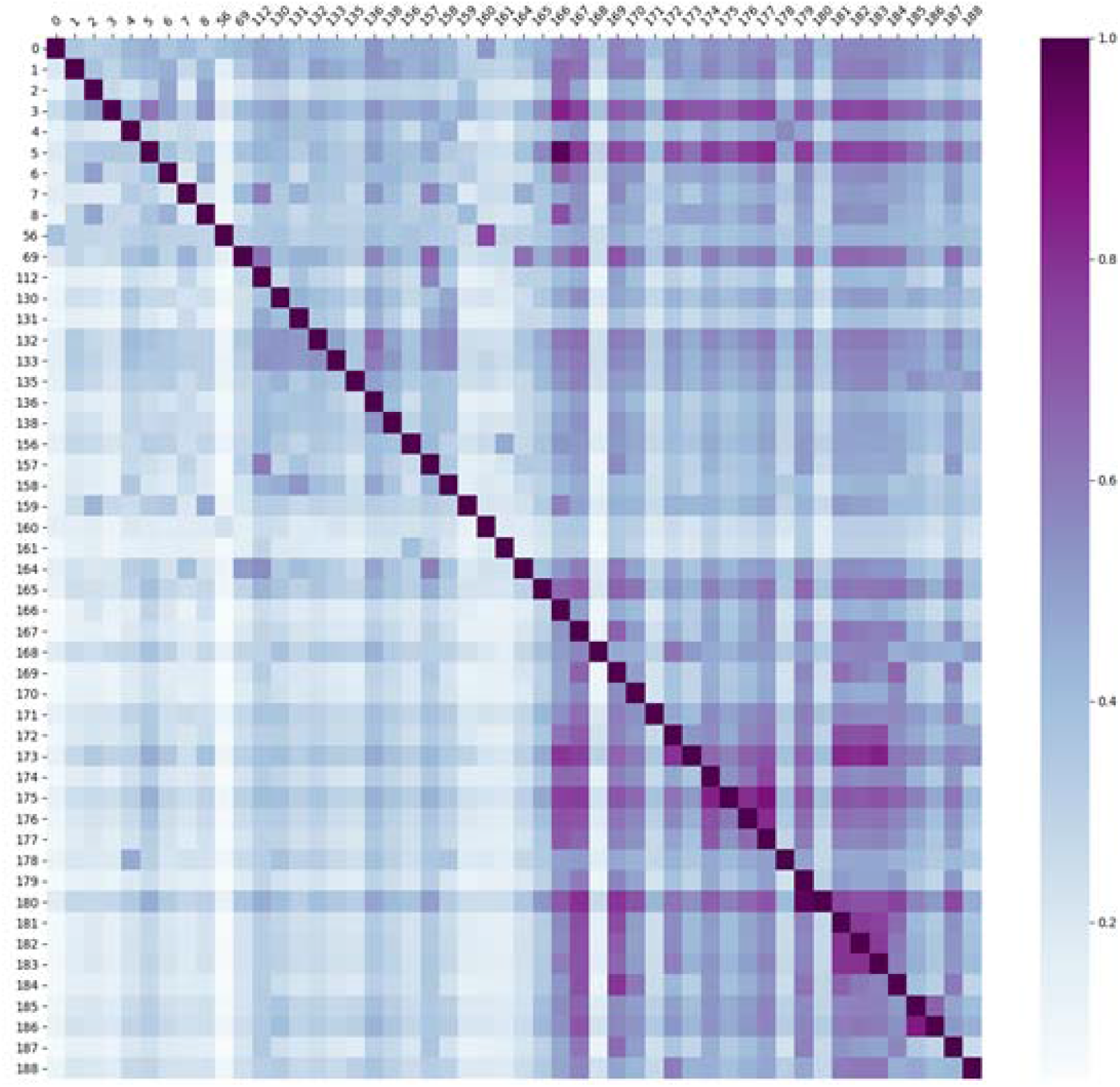
The correlations of active enhancers loci across different cell types. The result shows that the overall similarity between the 50 cell types is not very high, which highlights the importance of combining these cell types together as the training data. The cell type IDs and the name of the cell types can be referred to Table S2.

**Figure S3.**
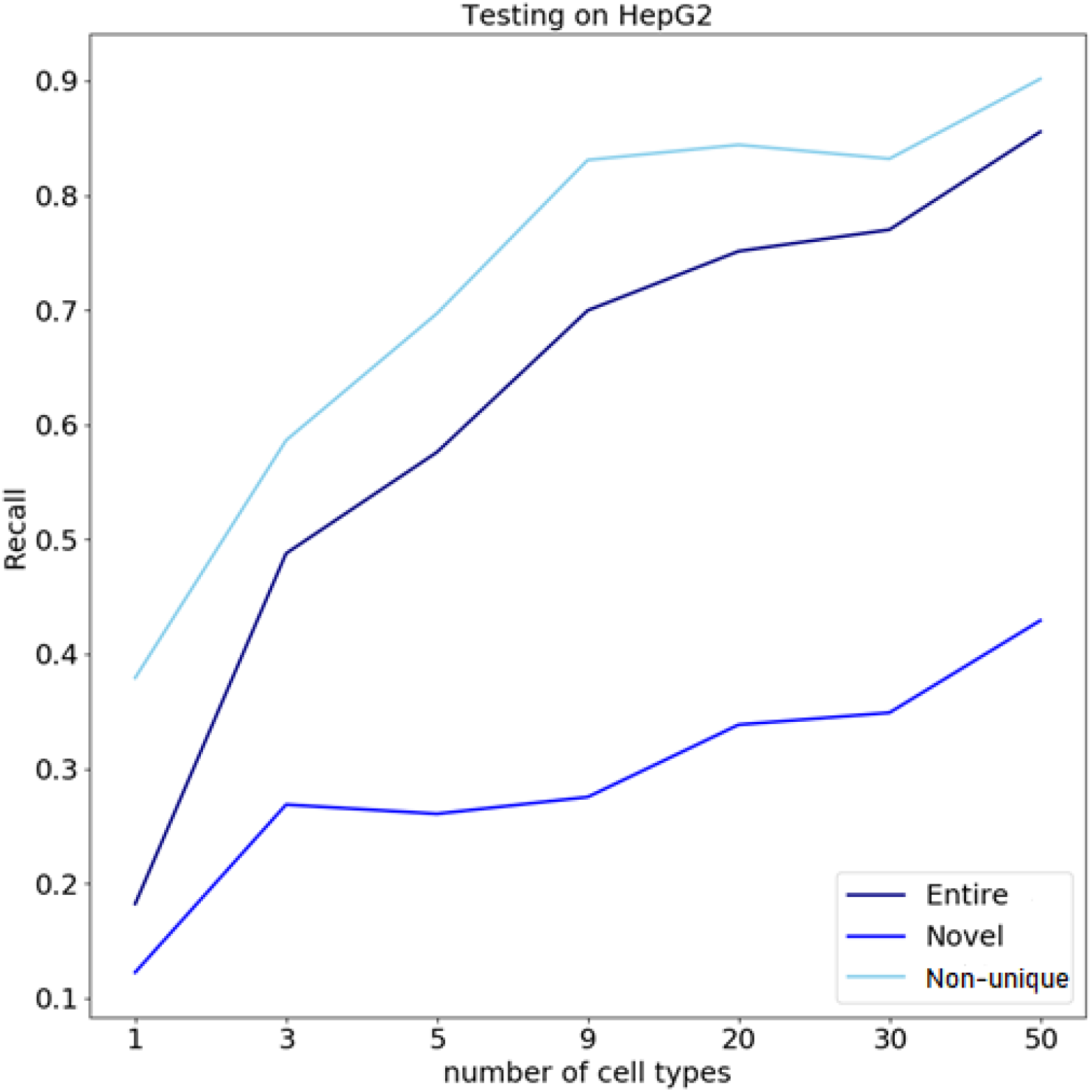
The recall rates of applying the models built with different numbers of cell types on the testing data (HepG2). The ‘Entire’ set contains all the instances in the testing data, while the set of ‘Novel’ denotes the instances that are active enhancers uniquely observed in HepG2 and the set of ‘Non-unique’ denotes the instances that have also been labeled as active enhancers by any of the cell types used in training. The figure shows that not only the recall of the non-unique active enhancers increases while including more cell types, but also the recall of novel active enhancers increases, indicating accuEnhancer can learn regulatory patterns more accurately when integrating more cell types in the training data.

**Figure S4.**
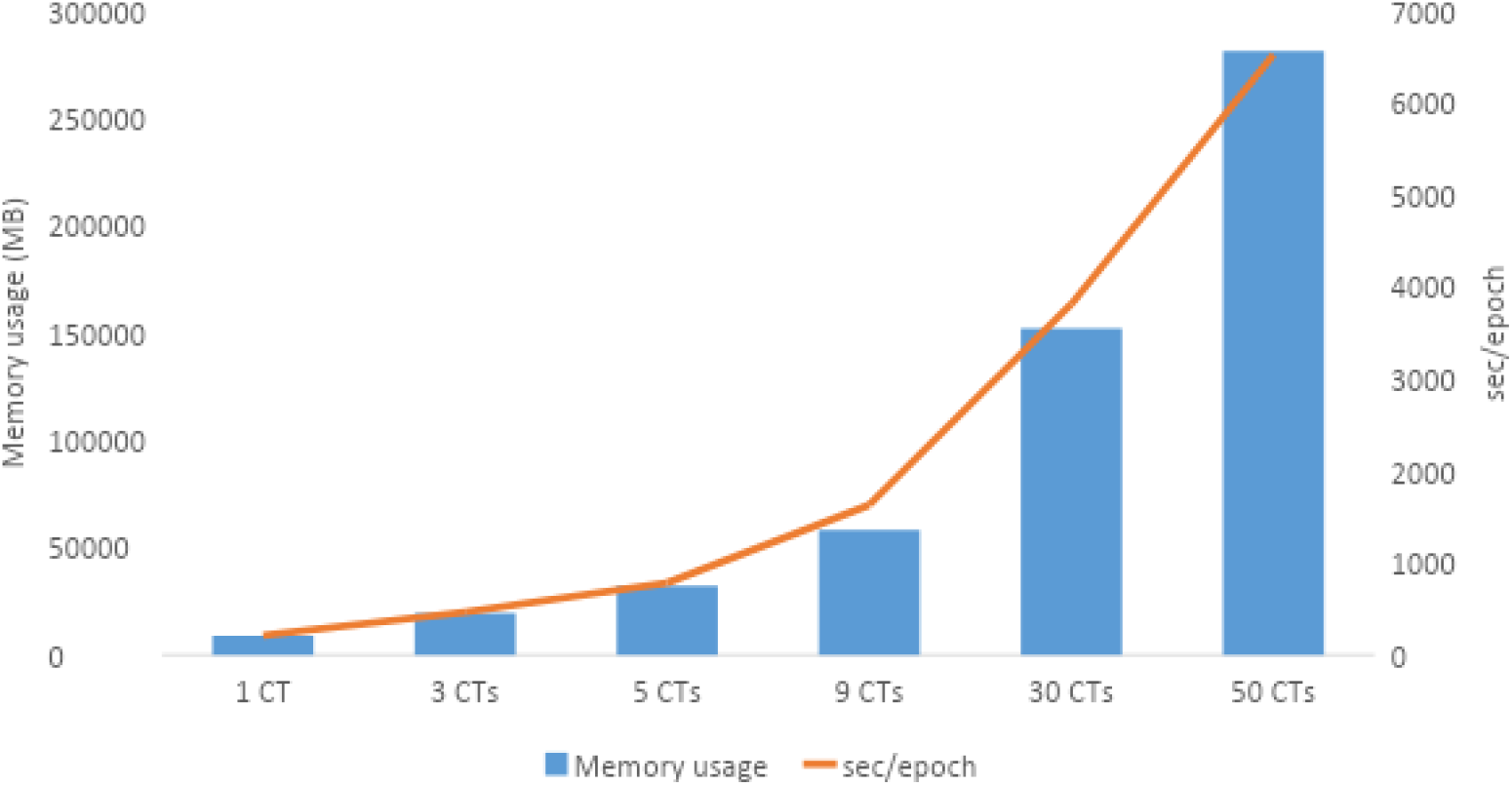
The resource usage of training accuEnhancer. The blue bar shows the peak memory usage during the total training process and the orange line shows the training duration of each epoch when different numbers of cell types are used in training. As the number of cell types used in the training data increases, the memory usage and the required training time also increase.

**Figure S5.**
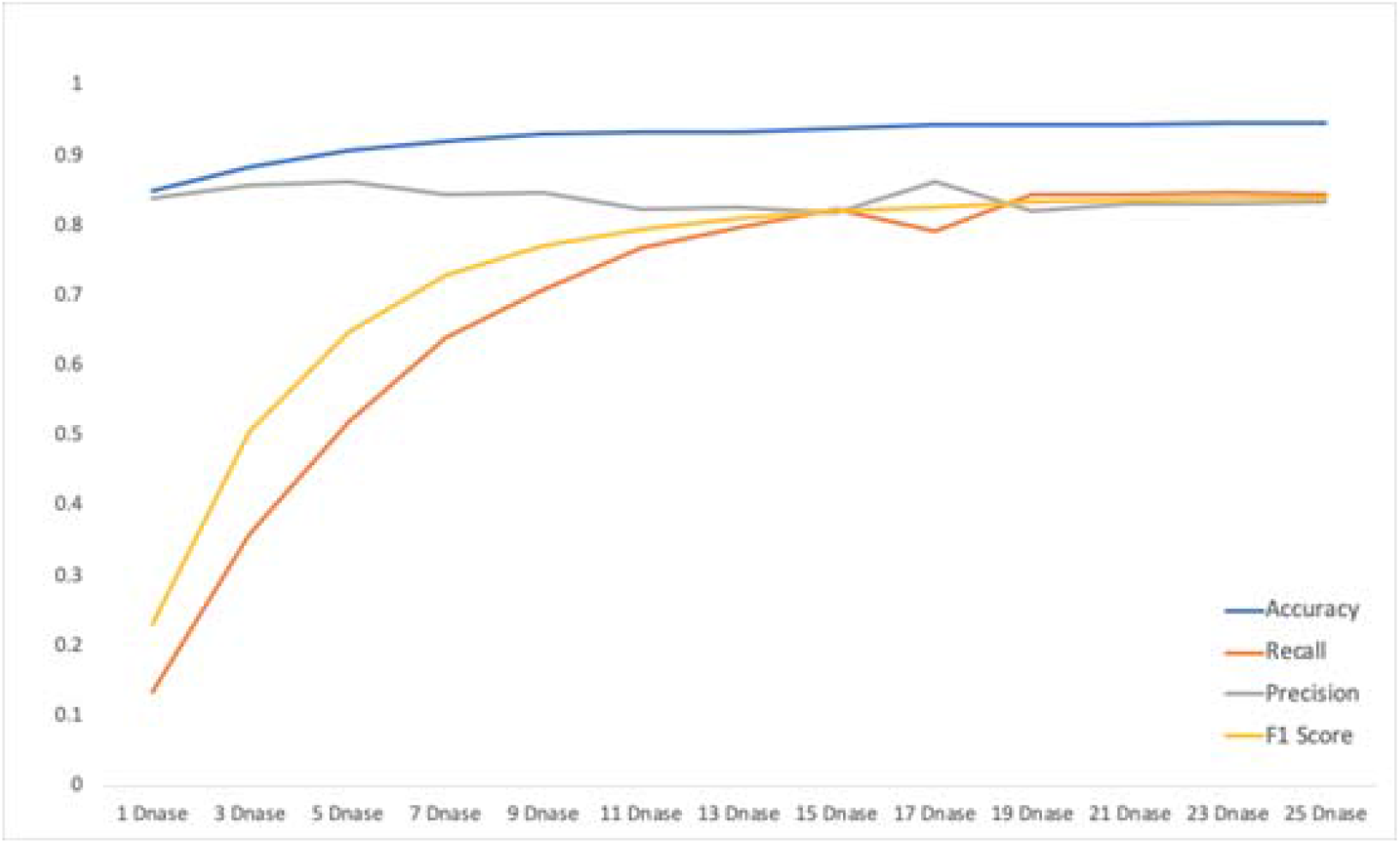
The testing (HepG2) performance of the models using different numbers of DNase bins in the functional data module. The training data included all the 50 cell types listed in Table S2.

**Figure S6.**
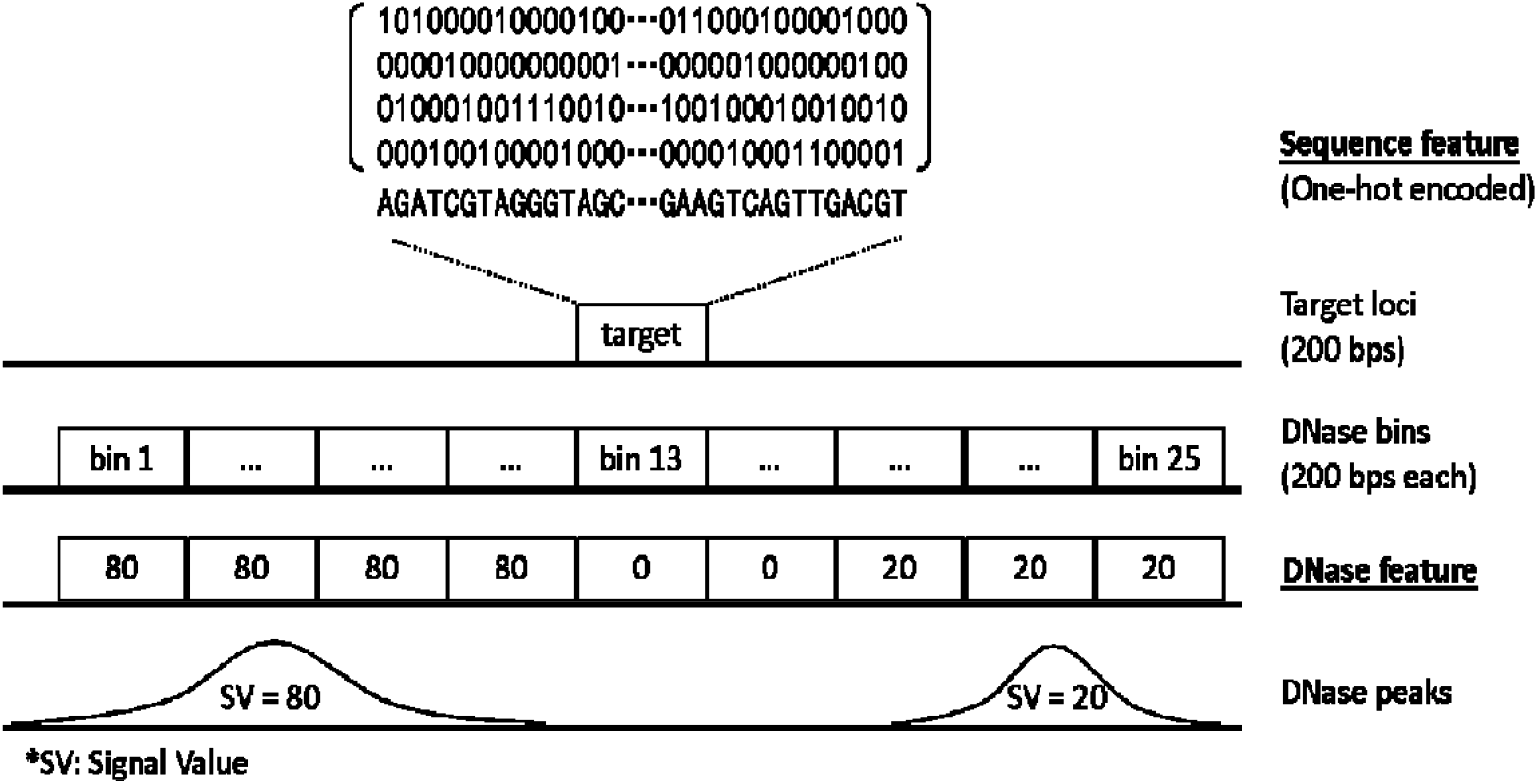
An overview of the input features for a given target training loci. The DNase signal values shown in the figure are the raw feature scores extracted from the ENCODE database narrow peak files. The signal value of the peak will be assigned if the bin is covered by the peak for more than 100 bps (50% of the bin length). Each target loci will contain 4×200 one-hot encoded sequence features and 25 DNase feature scores as describe in the Methods section.

**Table S1.** Cross-cell type prediction using a comprehensive set of functional data as the input features (H3K27me3, H3K36me3, H3K4me1, H3K4me2, H3K4me3, H3k9ac, DNase).

|  |  | Testing cell type |  |  |  |
| --- | --- | --- | --- | --- | --- |
| Training<br>cell type | F1 score | GM12878 | HepG2 | H1-hESC | K562 |
|  | GM12878 | 0.901 | 0.804 | 0.738 | 0.826 |
|  | HepG2 | 0.616 | 0.859 | 0.437 | 0.767 |
|  | H1-hESC | 0.875 | 0.801 | 0.813 | 0.801 |
|  | K562 | 0.852 | 0.817 | 0.701 | 0.882 |

**Table S2.** The list of 50 cell lines or tissues used in the training process. The narrow peak files of the DNase data and H3K27ac ChIP-seq data were downloaded from the ENCODE database.

| Cell type ID | Name | Cell type ID | Name |
| --- | --- | --- | --- |
| 0 | H1_BMP4_Derived_Mesendoderm_Cultured_Cells | 164 | AG04450 |
| 1 | K562_Leukemia_Cells | 165 | adrenal gland (51 yrs female) |
| 2 | Oci-Ly-7_B_Cell_Non-Hodgkin_Lymphoma_Cell_Line | 166 | spleen 54 yrs male |
| 3 | Thymus | 167 | stomach 51 yrs female |
| 4 | Chorion | 168 | small intestine 34 yrs male |
| 5 | Spleen | 169 | sigmoid colon female adult (51 year) |
| 6 | Karpas-422_B_Cell_Non-Hodgkin_Lymphoma_Cell_Line | 170 | gastrocnemius medialis female adult (51 year) |
| 7 | H1_Derived_Mesenchymal_Stem_Cells | 171 | thyroid gland male adult (54 years) |
| 8 | GM12878_Lymphoblastoid_Cells | 172 | female adult (53 years) Peyer's patch tissue |
| 56 | H9_Derived_Smooth_Muscle_Cultured_Cells | 173 | Peyer's patch male adult (54 years) |
| 69 | IMR90_Fetal_Lung_Fibroblasts_Cell_Line | 174 | upper lobe of left lung female adult (53 years) |
| 112 | SK-N-SH_Neuroblastoma_Cell_Line | 175 | upper lobe of left lung male adult (37 years) |
| 130 | MCF-7_Mammary_Gland_Adenocarcinoma_Cell_Line | 176 | upper lobe of left lung male adult (54 years) |
| 131 | PC-3_Prostate_Adenocarcinoma_Cell_Line | 177 | upper lobe of left lung female adult (51 year) |
| 132 | HeLa-S3_Cervical_Carcinoma_Cell_Line | 178 | placenta female embryo (113 days) |
| 133 | HCT116_Colorectal_Carcinoma_Cell_Line | 179 | tibial nerve male adult (37 years) |
| 135 | HepG2_Hepatocellular_Carcinoma_Cell_Line | 180 | tibial nerve female adult (51 year) |
| 136 | A549_EtOH_pt02pct_Lung_Carcinoma_Cell_Line | 181 | transverse colon female adult (53 years) |
| 138 | Panc1_Epithelioid_Carcinoma_Cell_Line | 182 | transverse colon female adult (51 year) |
| 156 | A673 | 183 | transverse colon male adult (37 years) |
| 157 | gm23248 | 184 | heart left ventricle female adult (53 years) |
| 158 | PC-9 | 185 | body of pancreas male adult (37 years) |
| 159 | MM1S | 186 | body of pancreas female adult (51 year) |
| 160 | gm23338 | 187 | tibial artery male adult (54 years) |
| 161 | SK-N-MC | 188 | large intestine male embryo (108 days) |

## References

Andersson, R., C. Gebhard, I. Miguel-Escalada, I. Hoof, J. Bornholdt, M. Boyd, Y. Chen, X. Zhao, C. Schmidl, T. Suzuki, E. Ntini, E. Arner, E. Valen, K. Li, L. Schwarzfischer, D. Glatz, J. Raithel, B. Lilje, N. Rapin, F. O. Bagger, M. Jørgensen, P. R. Andersen, N. Bertin, O. Rackham, A. M. Burroughs, J. K. Baillie, Y. Ishizu, Y. Shimizu, E. Furuhata, S. Maeda, Y. Negishi, C. J. Mungall, T. F. Meehan, T. Lassmann, M. Itoh, H. Kawaji, N. Kondo, J. Kawai, A. Lennartsson, C. O. Daub, P. Heutink, D. A. Hume, T. H. Jensen, H. Suzuki, Y. Hayashizaki, F. Müller, A. R. R. Forrest, P. Carninci, M. Rehli, A. Sandelin and F. C. The (2014). “An atlas of active enhancers across human cell types and tissues.” Nature 507(7493): 455–461.

Bernstein, B. E., J. A. Stamatoyannopoulos, J. F. Costello, B. Ren, A. Milosavljevic, A. Meissner, M. Kellis, M. A. Marra, A. L. Beaudet, J. R. Ecker, P. J. Farnham, M. Hirst, E. S. Lander, T. S. Mikkelsen and J. A. Thomson (2010). “The NIH Roadmap Epigenomics Mapping Consortium.” Nat Biotechnol 28(10): 1045–1048.

Bono, H., T. Kasukawa, M. Furuno, Y. Hayashizaki and Y. Okazaki (2002). “FANTOM DB: database of Functional Annotation of RIKEN Mouse cDNA Clones.” Nucleic Acids Res 30(1): 116–118.

Bu, H., Y. Gan, Y. Wang, S. Zhou and J. Guan (2017). “A new method for enhancer prediction based on deep belief network.” BMC Bioinformatics 18(12): 418.

Chen, K., Z. Chen, D. Wu, L. Zhang, X. Lin, J. Su, B. Rodriguez, Y. Xi, Z. Xia, X. Chen, X. Shi, Q. Wang and W. Li (2015). “Broad H3K4me3 is associated with increased transcription elongation and enhancer activity at tumor-suppressor genes.” Nature Genetics 47(10): 1149–1157.

Cirulli, E. T. and D. B. Goldstein (2010). “Uncovering the roles of rare variants in common disease through whole-genome sequencing.” Nature Reviews Genetics 11(6): 415–425.

Consortium, E. P. (2004). “The ENCODE (ENCyclopedia Of DNA Elements) Project.” Science 306(5696): 636–640.

Creyghton, M. P., A. W. Cheng, G. G. Welstead, T. Kooistra, B. W. Carey, E. J. Steine, J. Hanna, M. A. Lodato, G. M. Frampton, P. A. Sharp, L. A. Boyer, R. A. Young and R. Jaenisch (2010). “Histone H3K27ac separates active from poised enhancers and predicts developmental state.” Proc Natl Acad Sci U S A 107(50): 21931–21936.

Ernst, J. and M. Kellis (2012). “ChromHMM: automating chromatin-state discovery and characterization.” Nat Methods 9(3): 215–216.

Firpi, H. A., D. Ucar and K. Tan (2010). “Discover regulatory DNA elements using chromatin signatures and artificial neural network.” Bioinformatics 26(13): 1579–1586.

Griffiths-Jones, S., S. Moxon, M. Marshall, A. Khanna, S. R. Eddy and A. Bateman (2005). “Rfam: annotating non-coding RNAs in complete genomes.” Nucleic Acids Res 33(Database issue): D121–124.

He, Y., D. U. Gorkin, D. E. Dickel, J. R. Nery, R. G. Castanon, A. Y. Lee, Y. Shen, A. Visel, L. A. Pennacchio, B. Ren and J. R. Ecker (2017). “Improved regulatory element prediction based on tissue-specific local epigenomic signatures.” Proc Natl Acad Sci U S A 114(9): E1633–E1640.

Heintzman, N. D., R. K. Stuart, G. Hon, Y. Fu, C. W. Ching, R. D. Hawkins, L. O. Barrera, S. Van Calcar, C. Qu, K. A. Ching, W. Wang, Z. Weng, R. D. Green, G. E. Crawford and B. Ren (2007). “Distinct and predictive chromatin signatures of transcriptional promoters and enhancers in the human genome.” Nat Genet 39(3): 311–318.

Hoffman, M. M., O. J. Buske, J. Wang, Z. P. Weng, J. A. Bilmes and W. S. Noble (2012). “Unsupervised pattern discovery in human chromatin structure through genomic segmentation.” Nature Methods 9(5): 473–488.

Hon, G., B. Ren and W. Wang (2008). “ChromaSig: A Probabilistic Approach to Finding Common Chromatin Signatures in the Human Genome.” PLOS Computational Biology 4(10): e1000201.

Karmodiya, K., A. R. Krebs, M. Oulad-Abdelghani, H. Kimura and L. Tora (2012). “H3K9 and H3K14 acetylation co-occur at many gene regulatory elements, while H3K14ac marks a subset of inactive inducible promoters in mouse embryonic stem cells.” BMC Genomics 13(1): 424.

Kim, S. G., M. Harwani, A. Grama and S. Chaterji (2016). “EP-DNN: A Deep Neural Network-Based Global Enhancer Prediction Algorithm.” Scientific reports 6: 38433–38433.

Kleftogiannis, D., P. Kalnis and V. B. Bajic (2015). “DEEP: a general computational framework for predicting enhancers.” Nucleic Acids Res 43(1): e6.

Lawrence, S., C. L. Giles, A. C. Tsoi and A. D. Back (1997). “Face recognition: a convolutional neural-network approach.” IEEE Trans Neural Netw 8(1): 98–113.

Lee, D., R. Karchin and M. A. Beer (2011). “Discriminative prediction of mammalian enhancers from DNA sequence.” Genome Res 21(12): 2167–2180.

Liu, F., H. Li, C. Ren, X. Bo and W. Shu (2016). “PEDLA: predicting enhancers with a deep learning-based algorithmic framework.” Sci Rep 6: 28517.

Lonsdale, J., J. Thomas, M. Salvatore, R. Phillips, E. Lo, S. Shad, R. Hasz, G. Walters, F. Garcia, N. Young, B. Foster, M. Moser, E. Karasik, B. Gillard, K. Ramsey, S. Sullivan, J. Bridge, H. Magazine, J. Syron, J. Fleming, L. Siminoff, H. Traino, M. Mosavel, L. Barker, S. Jewell, D. Rohrer, D. Maxim, D. Filkins, P. Harbach, E. Cortadillo, B. Berghuis, L. Turner, E. Hudson, K. Feenstra, L. Sobin, J. Robb, P. Branton, G. Korzeniewski, C. Shive, D. Tabor, L. Q. Qi, K. Groch, S. Nampally, S. Buia, A. Zimmerman, A. Smith, R. Burges, K. Robinson, K. Valentino, D. Bradbury, M. Cosentino, N. Diaz-Mayoral, M. Kennedy, T. Engel, P. Williams, K. Erickson, K. Ardlie, W. Winckler, G. Getz, D. DeLuca, D. MacArthur, M. Kellis, A. Thomson, T. Young, E. Gelfand, M. Donovan, Y. Meng, G. Grant, D. Mash, Y. Marcus, M. Basile, J. Liu, J. Zhu, Z. D. Tu, N. J. Cox, D. L. Nicolae, E. R. Gamazon, H. K. Im, A. Konkashbaev, J. Pritchard, M. Stevens, T. Flutre, X. Q. Wen, E. T. Dermitzakis, T. Lappalainen, R. Guigo, J. Monlong, M. Sammeth, D. Koller, A. Battle, S. Mostafavi, M. McCarthy, M. Rivas, J. Maller, I. Rusyn, A. Nobel, F. Wright, A. Shabalin, M. Feolo, N. Sharopova, A. Sturcke, J. Paschal, J. M. Anderson, E. L. Wilder, L. K. Derr, E. D. Green, J. P. Struewing, G. Temple, S. Volpi, J. T. Boyer, E. J. Thomson, M. S. Guyer, C. Ng, A. Abdallah, D. Colantuoni, T. R. Insel, S. E. Koester, A. R. Little, P. K. Bender, T. Lehner, Y. Yao, C. C. Compton, J. B. Vaught, S. Sawyer, N. C. Lockhart, J. Demchok and H. F. Moore (2013). “The Genotype-Tissue Expression (GTEx) project.” Nature Genetics 45(6): 580–585.

Lu, Y., W. Qu, G. Shan and C. Zhang (2015). “DELTA: A Distal Enhancer Locating Tool Based on AdaBoost Algorithm and Shape Features of Chromatin Modifications.” PLOS ONE 10(6): e0130622.

Mardis, E. R. (2008). “The impact of next-generation sequencing technology on genetics.” Trends in genetics 24(3): 133–141.

Natoli, G. and J. C. Andrau (2012). “Noncoding Transcription at Enhancers: General Principles and Functional Models.” Annual Review of Genetics 46: 1–19.

Rajagopal, N., W. Xie, Y. Li, U. Wagner, W. Wang, J. Stamatoyannopoulos, J. Ernst, M. Kellis and B. Ren (2013). “RFECS: A Random-Forest Based Algorithm for Enhancer Identification from Chromatin State.” PLOS Computational Biology 9(3): e1002968.

Schwessinger, R., M. Gosden, D. Downes, R. C. Brown, A. M. Oudelaar, J. Telenius, Y. W. Teh, G. Lunter and J. R. Hughes (2020). “DeepC: predicting 3D genome folding using megabase-scale transfer learning.” Nature Methods.

Shlyueva, D., G. Stampfel and A. Stark (2014). “Transcriptional enhancers: from properties to genome-wide predictions.” Nat Rev Genet 15(4): 272–286.

Spitz, F. and E. E. Furlong (2012). “Transcription factors: from enhancer binding to developmental control.” Nat Rev Genet 13(9): 613–626.

Visel, A., S. Minovitsky, I. Dubchak and L. A. Pennacchio (2007). “VISTA Enhancer Browser--a database of tissue-specific human enhancers.” Nucleic Acids Res 35(Database issue): D88–92.

Wilken, M. S., J. A. Brzezinski, A. La Torre, K. Siebenthall, R. Thurman, P. Sabo, R. S. Sandstrom, J. Vierstra, T. K. Canfield, R. S. Hansen, M. A. Bender, J. Stamatoyannopoulos and T. A. Reh (2015). “DNase I hypersensitivity analysis of the mouse brain and retina identifies region-specific regulatory elements.” Epigenetics & Chromatin 8(1): 8.

Yang, B., F. Liu, C. Ren, Z. Ouyang, Z. Xie, X. Bo and W. Shu (2017). “BiRen: predicting enhancers with a deep-learning-based model using the DNA sequence alone.” Bioinformatics 33(13): 1930–1936.

Zhang, F. and J. R. Lupski (2015). “Non-coding genetic variants in human disease.” Hum Mol Genet 24(R1): R102–110.

Zhu, Y., L. Sun, Z. Chen, J. W. Whitaker, T. Wang and W. Wang (2013). “Predicting enhancer transcription and activity from chromatin modifications.” Nucleic Acids Res 41(22): 10032–10043.

